# Potential Energy Weighted Reactive Flux and Total Rate of Change of Potential Energy: Theory and Illustrative Applications

**DOI:** 10.1101/2022.07.08.499260

**Authors:** Wenjin Li

## Abstract

Reactive flux can be largely non-zero in a nonequilibrium ensemble of trajectories and provide insightful information for reactive transitions from the reactant state to the product state. Based on the reactive flux, a theoretical framework is proposed here for two quantities, the potential energy weighted reactive flux and the total rate of change of potential energy, which are useful for the identification of mechanism from a nonequilibrium ensemble. From such quantities, two multidimensional free energy analogues can be derived in the subspace of collective variables and they are equivalent in the regions where the reactive flux is divergence-free. These free energy analogues are assumed to be closely related to the free energy in the subspace of collective variables and they are reduced in the one-dimensional case to be the ensemble average of the potential energy weighted with reactive flux intensity, which was proposed recently and could be decomposed into energy components at the per-coordinate level. In the subspace of collective variables, the decomposition of the multidimensional free energy analogues at the per-coordinate level is theoretically possible and is numerically difficult to be calculated. Interestingly, the total rate of change of potential energy is able to identify the location of the transition state ensemble or the stochastic separatrix, in addition to the locations of the reactant and product states. The total rate of change of potential energy can be decomposed at the per-coordinate level and its components can quantify the contribution of a coordinate to the reactive transition in the subspace of collective variables. We then illustrated the main insights and objects that can be provided by the approach in the application to the alanine peptide in vacuum in various nonequilibrium ensembles of short trajectories and the results from these ensembles were found to be consistent.

## 1 INTRODUCTION

In the study of complex biomolecular systems with molecular dynamics (MD) simulations, one of the main challenges is the identification of mechanism.^1,2^ Rare transitions in biomolecular systems, such as large conformational changes of biomolecules, enzymatic reactions, and protein-ligand interactions, can be generally reduced to a prototypical problem of reactive transitions between two metastable states separated by an activation barrier,^3^ which are usually named as the reactant state and the product state in chemical reactions. In such a problem, the main task corresponds to identification of reaction coordinates from all the degrees of freedom of the system. The first step in such a task is usually the collection of all the reactive trajectories or transition paths that connect one metastable state to another, which forms the transition path ensemble (TPE) and is believed to contain all the information necessary for reaction coordinate identification. ^4–7^ The sampling of TPE is computational less expensive than the sampling of the equilibrium ensemble and can be accomplished by various path-sampling approaches, such as transition path sampling (TPS),^4,8–11^ nudged elastic-band method,^12,13^ finite-temperature string method, ^14,15^ and forward flux sampling. ^16,17^

However, reaction coordinate identification from the TPE is only possible if an approach to analyse the transition paths is available. In the early attempts to this challenge, many approaches require the information of committor or splitting probability. The committor is the probability to commit to the product state first when simulations initiate from a given configuration with momenta assigned from a Boltzmann distribution ^18–20^ and is assumed to be an ‘ideal’ one-dimensional representation of reaction coordinates.^7^ For a diffusive process with a parabolic barrier, a one-dimensional representation of reaction coordinates can be calculated from the committor and it is the eigenvector of the only positive eigenvalue of the matrix (−***HD***),^21,22^ where ***H*** is the Hessian matrix of the potential energy and ***D*** is the diffusion coefficient matrix. Pioneered approaches in the identification of reaction coordinate from the TPE are genetic neural network method^23^ and likelihood maximization method,^24–26^ which select reaction coordinates out of a pool of candidate coordinates with the committor as the reference. We refer to a recent review^7^ for the details and other early works in such an endeavour.

Recent efforts to identify reaction coordinates from the TPE focused on the development of approaches that did not require extra simulations and thus avoided the need to compute a very large number of trajectories for committor calculation. Li used the equipartition terms in the TPE along a coordinate to appraise the relevance of the coordinate to reaction coordinates and the maximization of equipartition terms reproduced a good one-dimensional representation of reaction coordinates.^27^ Later on, a density-weighted average of the flux along a coordinate over the TPE was demonstrated to be a promising quantity to rank reaction coordinates^28^ and its relation to the transition path theory, a statistical theory for describing the TPE pioneered by Weinan and Vanden-Eijnden,^29^ was clarified thereafter.^30^ On the other hand, the committor can be efficiently estimated by methods other than shooting many unconstrained trajectories with random assignment of momenta and then committor-based approaches can be applied to identify reaction coordinates. While the committor of configurations in the TPE can be quickly evaluated by a fitting procedure, ^31^ the committor of configurations in non-equilibrium ensembles of trajectories other than the TPE can be estimated by such algorithms as non-equilibrium non-parametric analysis, ^32^ iso-committor surfaces calculation, ^33^ and Markov state models.^34,35^ Without the information of committor, machine learning methods were recently employed to identify reaction coordinates for biomolecular systems as well. ^36,37^

In the above-mentioned approaches for reaction coordinate identification from the TPE, the importance of a coordinate to a rare transition is quantified by a single value, which could be the equipartition term of a coordinate, ^27^ the density-weighted average of the flux along a coordinate, the variation of a coordinate along the stochastic separatrix in transition state ensemble optimisation, ^6,38^ or the root-mean-square error in genetic neural network method.^23^ However, the contribution of a coordinate to a transition can varies as the transition progresses. As a hypothetical example, a coordinate may play an important role in climbing up the energy barrier, while its role is negligible when rolling down the barrier. Thus, a single value of a quantity can just quantify the averaged contribution of a coordinate to a transition process and it should be more informative if one can quantify the contribution of a coordinate at various stages of a transition. Since the reaction coordinates can quantify the progression of a transition, it is thus a good idea to estimate the importance of a coordinate as a function of reaction coordinates. As a first attempt towards the goal, Li and co-workers proposed the emergent potential energy on a coordinate induced by a one-dimensional representation of reaction coordinates, which is a function of the one-dimensional reaction coordinate, to appraise the relevance of the coordinate to reaction coordinates during the whole transition process.^39^ However, the physical meaning of the emergent potential energy was not very clear until it was expressed as an ensemble average weighted with flux intensity and its connection was made to the theoretical framework of energy decomposition along reaction coordinate (EDARC).^40^ In the EDARC approach, a free energy analogue, which is a close approximation of the free energy, along a one-dimensional reaction coordinate was decomposed into energy components on each coordinate. The component on a coordinate along the one-dimensional reaction coordinate, thus a function of such a reaction coordinate, was demonstrated to be capable of quantifying its contribution to a transition process at any point along the one-dimensional reaction coordinate. ^40^ Although both the emergent potential and the EDARC approach can quantify the relative importance of a coordinate in a transition process, the EDARC approach has a advantage over the emergent potential energy: The energy component on a coordinate in the EDARC approach quantify the absolute importance of a coordinate in unit of energy, as the summation of the energy components on all coordinates gives a free energy analogue that is very close to the free energy.

Although a one-dimensional reaction coordinate may be adequate to describe the progression of a transition very well in many systems, it can be very challenging to obtain such a one-dimensional reaction coordinate and thus often-times it requires several collective variables (CVs) to capture the effective dynamics of a complex system. When a set of CVs is provided, can one quantify the contribution of a coordinate at any point in the subspace of collective variables (CVs)? That is to say, can we extend the EDARC approach to multi-dimensional cases? Here, we will show that potential energy weighted reactive flux is a promising answer to these questions. The projection of potential energy weighted reactive flux onto the subspace of CVs results a quantity that defines a multi-dimensional free energy analogue, which is assumed to be a close approximation of the free energy in the subspace of CVs. The decomposition of such a multi-dimensional free energy analogue at the per-coordinate level provides a way to quantify the importance of a coordinate to a transition process in the subspace of CVs and it is the generalization of the EDARC approach to the multi-dimensional subspace of CVs. In the following, the theoretical framework is first introduced for potential energy weighted reactive flux and its close relative, i.e., the total rate of change of potential energy. Then, we illustrate the applications of such a theoretical framework in the *C*_7eq_ → *C*_7ax_ isomerization of the alanine dipeptide in vacuum, a system that has been intensively studied previously. ^23,27,28,30,39,41–43^ The viability of this approach is tested on several non-equilibrium ensembles of short trajectories that we have examined previously using EDARC.^40^ Finally, some concluding remarks are given at the end of the article.

## 2 Methods

### 2.1 Theory

#### 2.1.1 Flux and current

Let ***x***(*t*) representing an infinitely long equilibrium trajectory of a system with ***x*** being its positional coordinates. In an nonequilibrium ensemble *U* of trajectories, whose elements are taken from the equilibrium trajectory, and a generalized coordinate *ξ*(***x***), the flux passing through a dividing surface of *ξ*(***x***) = *ξ*′ according to Ref. [^30^] is given by

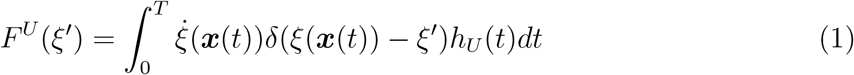

Here, *δ*(*x*) is the Dirac delta function and *h*_*U*_ (*t*) is an indicator function for the ensemble *U*. Specifically, *h*_*U*_ (*t*) = 1 if *t* ∈ *U* [i.e., at time *t*, ***x***(*t*) belongs to a trajectory in the nonequilibrium ensemble *U*] and *h*_*U*_ (*t*) = 0 otherwise. Note that, *F*^*U*^(*ξ*) is a function of the coordinate *ξ* and we call it the flux along the coordinate *ξ*. Usually, *ξ*(***x***) is chosen to be good one-dimensional representation of reaction coordinates. A particular choice of a nonequilibrium ensemble *U* could be the TPE, which consists of all the reactive trajectories taken from the long trajectory. For a system that have two metastable states A and B, a reactive trajectory from A to B is a piece of the equilibrium trajectory that starts from the state A and go to the state B without visiting the state A again. In the TPE, the flux passing through any isosurface of the reaction coordinate equals the number of reactive trajectories. ^30^ Thus, the flux along the reaction coordinate in the TPE is a constant and its derivative is zero. In a non-equilibrium steady state from A to B, the flux along the reaction coordinate is a constant as well and it equals the one in the corresponding TPE.

The flux or current in a subspace of CVs was also given in Ref. [^30^] and it is a vector field. Given a set of CVs ***y*** ≡ {*y*_1_, *y*_2_, …, *y*_*n*_}, whose elements are functions of ***x***, the current in the space of these CVs (we use flux when referring to a scalar and current when a vector field is referred from now on) is then defined as

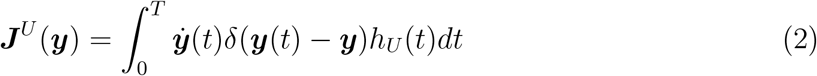

When *U* corresponds to the TPE or the nonequilirbirum steady state, the current in a subspace of reaction coordinates is divergences free, that is ∇ · ***J***^*U*^(***y***) = 0 when ***y*** can describe the effective dynamics from A to B of a system.

#### 2.1.2 Potential energy weighted current and total rate of change of potential energy

Previously we showed that the average energy weighted with flux along a one-dimensional representation of reaction coordinates defines a free energy analogue that is in close relationship with the free energy along the coordinate. ^40^ Following this path of thinking, it could be insightful in the configurational space to look at a potential energy weighted current for a nonequilibrium ensemble *U*, which is defined as

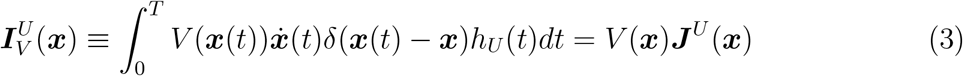

One immediately has,

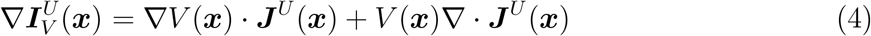

In case that the current in *U* is divergence free, i.e., ∇ · ***J***^*U*^(***x***) = 0 (e.g., *U* corresponds to the non-equilibrium steady state or the TPE), then

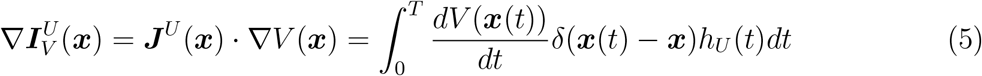

Thus, the divergence of the potential energy weighted current equals the directional derivative of the potential energy in the direction of the reactive current in case of a divergence-free current. We then introduce a quantity 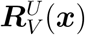,

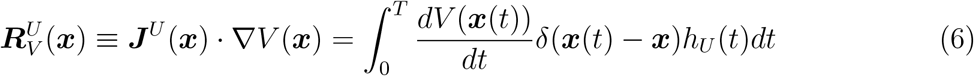

Actually, 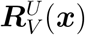 is the total rate of change of potential energy (TRCPE) at a point in the configurational space. It is the dot product between the reactive current and the gradient of potential energy and can be also viewed as the flux intensity along the gradient of the potential energy and is weighted by the magnitude of the gradient, i.e., 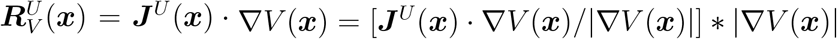.

The projection of 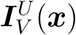 to the subspace of CVs gives (Appendix A),

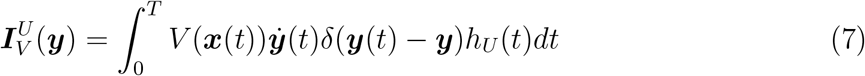

We name 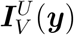 as the potential energy weighted current in the CVs subspace. Note that 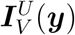 is a function of ***y*** and its dimension is a product of the dimension of energy and the one of the current ***J***^*U*^(***y***). We thus hypothesize that there exists an energy function *Ă*^*U*^(***y***) of ***y*** (thus a multidimensional free energy analogue in the subspace of CVs) such that 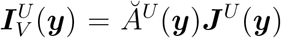. If the magnitude of the current ***J***^*U*^(***y***) is non-zero everywhere, one immediately finds that

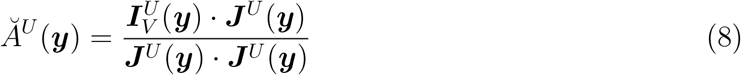

Thus, the projection of 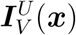 to the subspace of CVs defines a multidimensional free energy analogue *Ă*^*U*^(***y***). If only a one-dimensional representation of reaction coordinates *ξ* is given, i.e., the vector ***y*** is replaced by a scalar *ξ*, then we have,

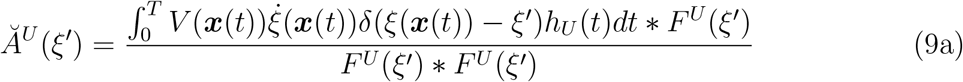

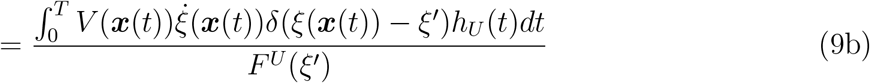

and the free energy analogue *Ă*^*U*^(*ξ*) in the one-dimensional case proposed previously^40^ is reproduced.

In analogue to the derivation of 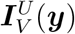 the projection of 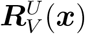 to the subspace of CVs gives,

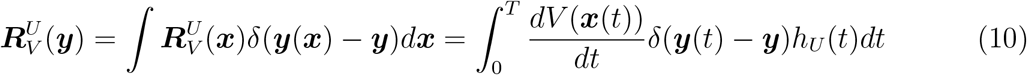

Thus, 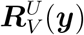 is the TRCPE at a point in the subspace of CVs. Since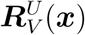 is the dot product between the reactive current and the gradient of potential energy, we also hypothesize that 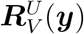 is the dot product between the reactive current and the gradient of an energy function in the subspace of CVs, i.e., there exists an energy function *Ã*^*U*^(***y***) of ***y*** (another multidimensional free energy analogue in the subspace of CVs) such that 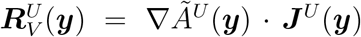. Then 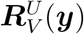 can be viewed as the flux intensity along the gradient of *Ã*^*U*^(***y***) and is weighted by the magnitude of the gradient, i.e., 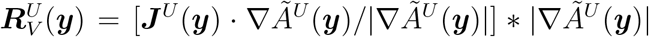 It is beyond our knowledge in mathematics to answer whether or not *Ã*^*U*^(***y***) is unique when 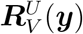 and ***J***^*U*^(***y***) are given. Here, we assume the uniqueness of *Ã*^*U*^(***y***) as *Ã*^*U*^(***y***) should be continuous and differentiable. Note that the potential energy function *V* (***x***) is usually a continuous and differentiable function. Under the assumption of a unique *Ã*^*U*^(***y***), we would like to point out that the numerical estimation of *Ã*^*U*^(***y***) is not trivial in case that 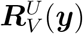 and ***J***^*U*^(***y***) are provided. It is easy to show that in the one-dimensional case, where a one-dimensional representation of reaction coordinates *ξ* is given,

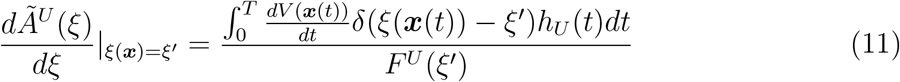

and the other free energy analogue *Ã*^*U*^(*ξ*) in the one-dimensional case proposed previously^40^ is reproduced as well.

In summary, two multidimensional free energy analogues *Ă*^*U*^(***y***) and *Ã*^*U*^(***y***) are defined from the potential energy weighted reactive current and they are the generalizations of the one-dimensional free energy analogues *Ă*^*U*^(*ξ*) and *Ã*^*U*^(*ξ*) to the multi-dimensional case, respectively.

In the region of the subspace of CVs where the divergence of the reactive current vanishes, i.e., there is no source or sink in this region for the ensemble *U*, we find that (Appendix B)

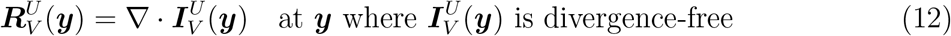

Under the assumption of a unique *Ã*^*U*^(***y***), it is obvious that *Ă*^*U*^(***y***) and *Ã*^*U*^(***y***) are equivalent in case of a divergence-free current.

#### 2.1.3 Decomposition of 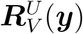

In the one-dimensional case, the free energy analogue *Ã*^*U*^(*ξ*) can be decomposed at the per-coordinate level. In the following, we will show that the multi-dimensional free energy analogue *Ã*^*U*^(***y***) can be decomposed into components for each coordinate as well. Let ***z*** ≡ {*z*_1_, *z*_2_, …, *z*_n_} being an independent and complete set of configurational coordinates, we have 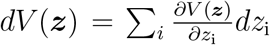 (the total differential of *V* (***z***)). Then, 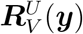 in eq. 10 can be rewritten as,

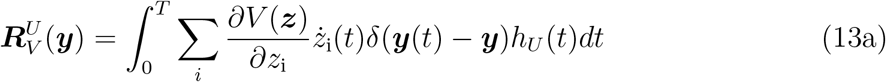

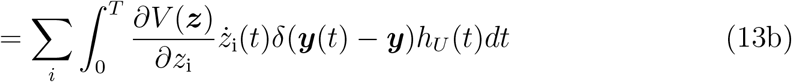

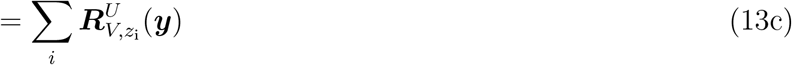

As indicated in eq. 13c, 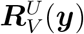 is now decomposed into components 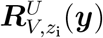 on each coordinate *z*_i_, with

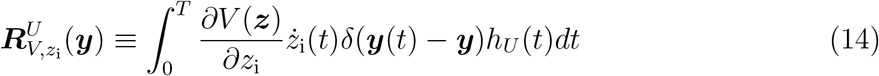

Thus, eq. 13c defines a decomposition of 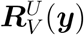 at the per-coordinate level. Now, we assume there exists a unique energy function 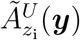 such that 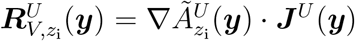 It is easy to show that 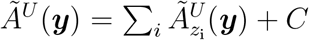 with *C* being a constant. Therefore, *Ã*^*U*^(***y***) can be decomposed at the per-coordinate level and 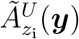 is its energy component on a coordinate *z*_i_. One would suggest that 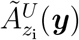 can be used to quantify the contribution of a coordinate *z*_i_ at any location of the subspace of CVs. Since the calculation of 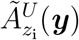 is mathematically as difficult as the estimation of *Ã*^*U*^(***y***), in the article we will decompose 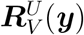 instead.

#### 2.1.4 Evaluation of 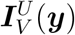 and 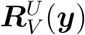 for complex systems

For complex systems, it is not practical to explore the energetics of the whole system at the per-coordinate level. Usually, a proper subsystem is selected to reduce the dimension and the noisiness due to the large fluctuations in the environment or bath modes. For example, it is shown to be feasible and useful when only the residues close to the retinal were included in the analysis of residue-residue mutual works in rhodopsin.^44^ Let us define that the subsystem is described by ***y***, and the rest of the system is described by ***y***′ (i.e., {***y, y***′} is another complete and independent set of configurational coordinates). In terms of {***y, y***′}, the potential energy *V* (***y, y***′) = *V*_***y***_(***y***) + *V*_***y***′_ (***y***′) + *V*_coupl_(***y, y***′). *V*_***y***_(***y***) and *V*_***y***′_ (***y***′) are the potential energy for the subsystem and its environment, respectively. *V*_coupl_(***y, y***′) is the interaction energy between the subsystem and its environment. We thus also assume that reaction coordinates are functions of ***y***, i.e., the set of CVs ***y*** can capture the essential dynamics of the transition from A to B, and ***y***′ are bath modes, whose behaviours in the ensemble *U* resemble the ones in an equilibrium ensemble.

The potential energy weighted current in the subspace of ***y*** can then be expressed as

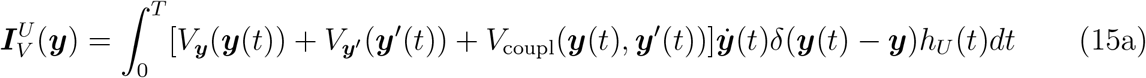

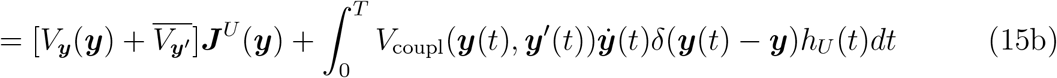

where, 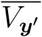 is the average of *V*_***y***′_ (***y***′) in the equilibrium ensemble and thus a constant, which can be ignored. Thus, by ignoring the term *V*_***y***′_ (***y***′), which is generally large and very noisy, the numerical estimation of 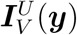 can be greatly improved.

On the other hand, we can write 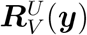 as

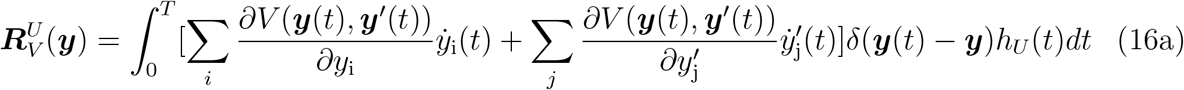

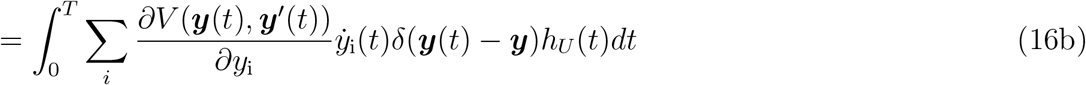

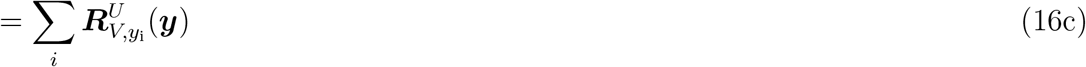

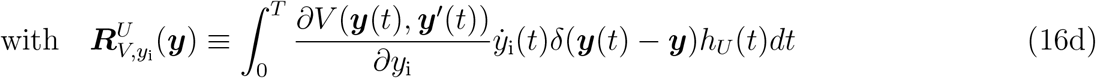

Here, a term 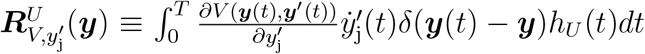 is the component of 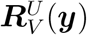 on a coordinate 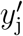 and it vanishes as the behaviour of 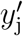 resembles the one in an equilibrium ensemble by assumption. Thus, 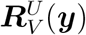 can be approximated by the summation of components on each coordinate in ***y*** and only the generalized forces on ***y*** are required for the calculation of 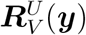. Usually, the number of variables in ***y*** is much smaller than the number of degrees of freedom of the system, and the computational cost is tremendously reduced by circumventing the need for the calculation of the generalized forces on bath modes.

From eq. 16, if the current in the subspace of CVs is non-zero, 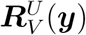 can be further rewritten as,

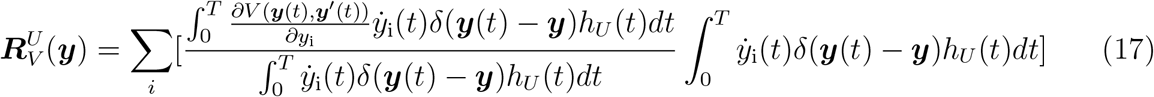

Note that 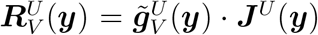 where 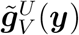 represents the gradient of the free energy analogue *Ã*^*U*^(***y***) to simplify the notation, i.e., 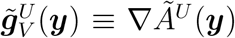. One natural solution for 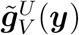 is given below with the *i*-th element of 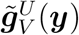 being provided by

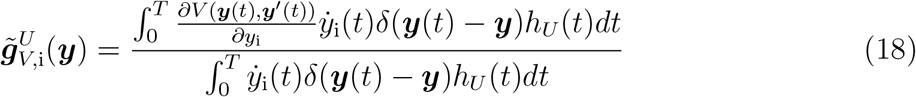

This equation provides a way to estimate *Ã*^*U*^(***y***), although the calculation of a multidimensional function from its gradient is numerically non-trivial.

##### Linear Coupling between the subsystem and its environment

In the following, we consider a special case that the coupling potential between the subsystem and its environment can be expressed as the product of a function of the subsystem and a function of its environment, i.e., *V*_coupl_(***y, y***′) = *g*(***y***)*m*(***y***′). Substituting *V* (***y, y***′) with *V*_***y***_(***y***) + *V*_***y***′_ (***y***′) + *V*_coupl_(***y, y***′) in eq. 18, we find that

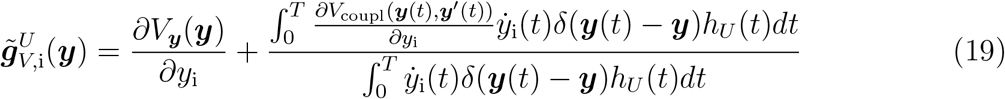

Since 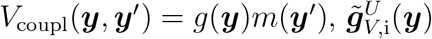 can then be written as

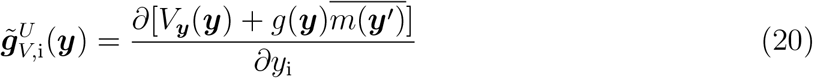

where, 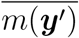 is the average of *m*(***y***′) in the equilibrium ensemble and thus a constant. Similarly, we find from eq. 15b that

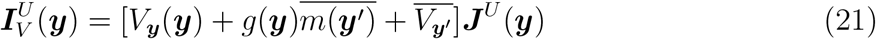

From eqs. 8 and 21, we have

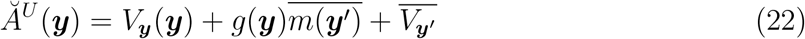

Note that 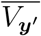 is a constant. Thus, under the assumption that the distribution of ***y***′ at any position of ***y*** resemble the one in the equilibrium ensemble, *Ã*^*U*^(***y***) is equivalent to *Ă*^*U*^(***y***) (i.e., differ by a constant). Obviously, when *V*_coupl_(***y***, ***y***′) can be expressed as the summation of functions with the same form of *g*(***y***)*m*(***y***′), this statement holds as well.

##### Weak coupling between the subsystem and its environment

If the coupling between the subsystem and its environment is weak, that is 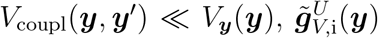 and *Ă*^*U*^(***y***) can be further approximated as

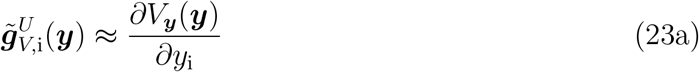

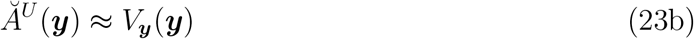

These results suggests that *Ă*^*U*^(***y***) and *Ã*^*U*^(***y***) are equal and both can be approximated by the potential energy of the subsystem under the assumption of weak coupling between the subsystem and its environment. Thus, the above derivations suggest that the free energy analogue and its energy components can be obtained from a proper subsystem that is a close approximation of the whole system. If the ∇*Ã*^*U*^(***y***) obtained with eq. 23a is close to the one with eq. 19, the subsystem could be considered a proper one that preserves the free energy analogue of the system. The energy components obtained from this subsystem are thus good approximations of the ones from the entire system.

#### 2.1.5 Overdamped dynamics and parabolic energy barrier

To explore the properties of 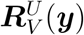 we consider two particular cases of the dynamics of a system: (1) Overdamped dynamics and parabolic energy barrier; (2) A slow reaction coordinate coupled with fast bath modes. First, we assume an overdamped dynamics and a parabolic barrier of the potential of mean force *A*(***y***) with the barrier top at ***y*** = 0, that is *βA*(***y***) = ***y***^*t*^***Hy****/*2, where *β* = 1*/*(*k*_B_*T*) and ***H*** is the Hessian matrix. From Roux^45^ in case of a position-independent diffusion matrix ***D***, the committor near the saddle point can be approximated by *q*(***y***) ≈ *q*(***y*** · ***e***) with ***e*** being the unit vector parallel to ∇*q*(***y***), and ***e*** is the eigenvector of the only negative eigenvalue *λ* of ***HD***, i.e., ***HDe*** = −*λ****e***. Since the reactive current in overdamped dynamics can be expressed as ^46^

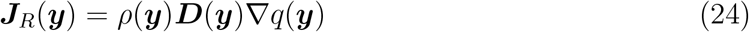

where *ρ*(***x***) is the equilibrium probability density in canonical ensemble and it is proportional to *e*^−*βA*(***y***)^. Then, we can write 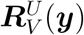 as

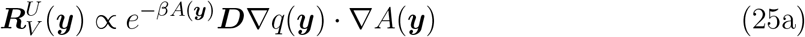

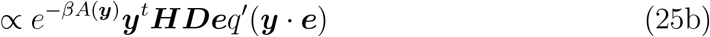

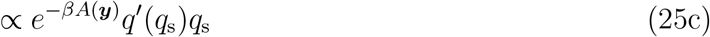

here, *q*_s_ = ***y*** · ***e*** is the reaction coordinate. In the stochastic separatrix or the transition state ensembles, i.e., *q*_s_ = 0, we have 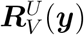 vanishes. Thus, the hypersurface of 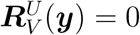 near the barrier top corresponds to the stochastic separatrix.

One may check the gradient of 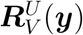 which can be expressed as

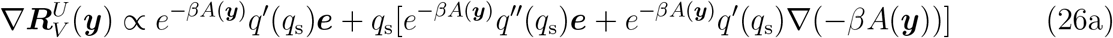

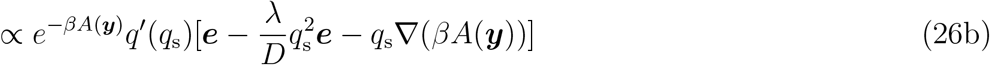

where, the conclusion that *q*^*′′*^(*q*_s_) = −*λq*_s_*q*^*′*^(*q*_s_)*/D* from Ref. [^45^] is used with *D* ≡ ***e***^*t*^***De***. The vector 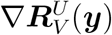 is thus found to be parallel to ***e*** in the vicinity of the separatrix, i.e., *q*_s_ ≈ 0.

#### 2.1.6 A slow reaction coordinate coupled with fast bath modes

Now we consider a system that consists of a slow coordinate *q*_s_, which is also the only reaction coordinate, and a pool of fast bath modes ***q***_b_. It is assumed that the dynamics along bath modes in a non-equilibrium ensemble *U* resembles the one in equilibrium and the coupling between the bath modes and the reaction coordinate *q*_s_ is weak. Since the components in the bath modes are vanishingly small, we can approximate 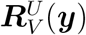 with

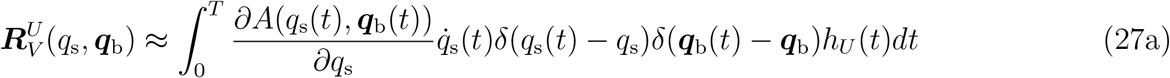

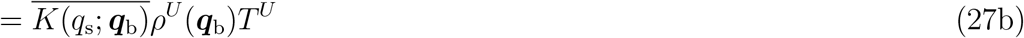

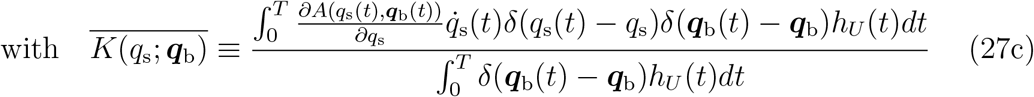

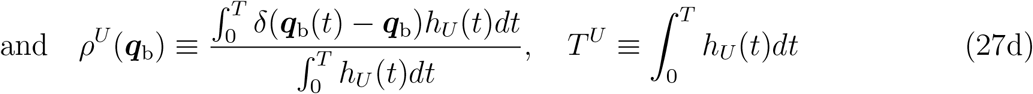

If 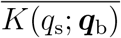 is independent to ***q***_b_, then

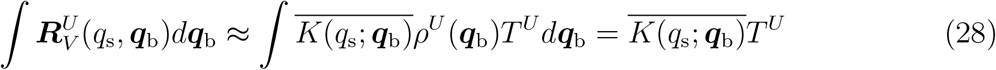

On the other hand,

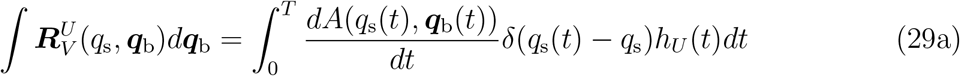

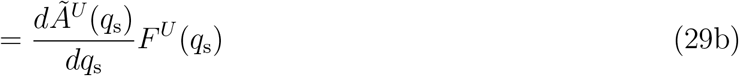

Thus, 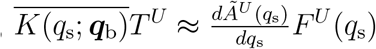 and

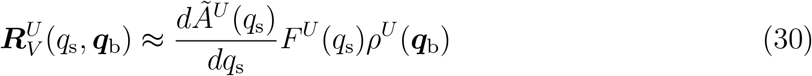

The gradient of 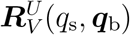 is

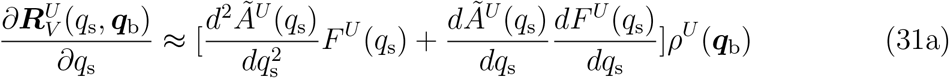

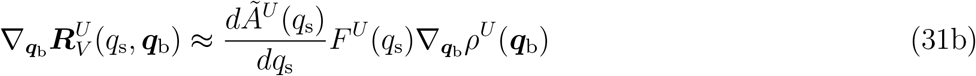

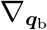 denotes the derivatives with respect to ***q***_b_. eq. 31 suggests that the gradient of 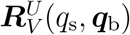 points the direction of the reaction coordinate when (1) 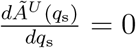 or (2) 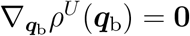, a zero vector.

One particular case when 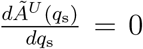 is at the barrier top of the free energy analogue (*Ã*^*U*^(*q*_s_)) along the reaction coordinate. It is the hypersurface 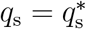 with 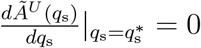 which could be the stochastic separatrix. At the hypersurface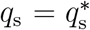, the directions of all 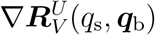 are parallel to the direction of the reaction coordinate *q*_s_ and the magnitude of 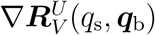 is proportional to *ρ*^*U*^(***q***_b_). The point at which the magnitude of 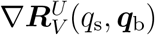 reaches its maximum corresponds to the saddle point. From eq. 30, when 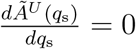, we have 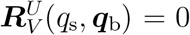. Thus, the hypersurface of 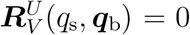 near the barrier top corresponds again to the stochastic separatrix, from which the reaction coordinate can be identified. It is obvious that 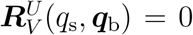 as well at the reactant state and the product state. Thus, the hypersurfaces of 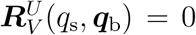 can be used to identify such important states as the transition state, the reactant state, and the product state.

A special case when 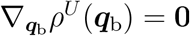 is a curve where *ρ*^*U*^(***q***_b_) reaches its maximum, which corresponds to the minimum free energy path and passes the saddle point. On the other hand, 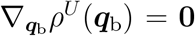 corresponds to the maximum of 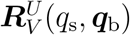 on the isosurfaces of the reaction coordinate *q*_s_, as indicated in eq. 30. Thus, the minimum free energy path and the direction of the reaction coordinate along it can be obtained via visual inspection of 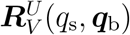 and 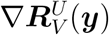 in the subspace of CVs, respectively.

#### 2.1.7 Generalization of potential energy weighted current

So far, the potential energy is used as the weight for the reactive current. Actually, an arbitrary function *f* (***x***) of ***x*** can be used as the weight and we can considered the reactive current weighted with *f* (***x***) in an ensemble *U* in the subspace of CVs,

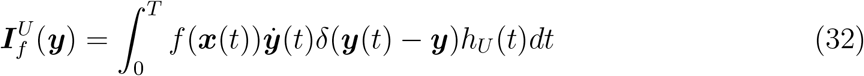

The corresponding expectation 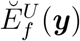 of *f* (***x***(*t*)) in the subspace of CVs is given by 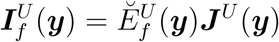. Similarly, the corresponding total rate of change of *f* (***x***) is,

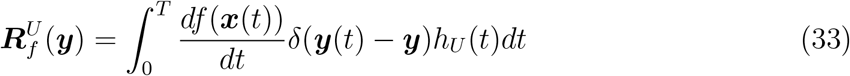

and another expectation 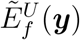 that satisfies 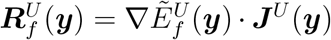 can also be defined. It is easy to show that 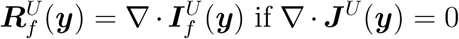. Then, 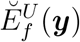 and 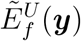 equals if one assumes the uniqueness of 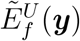 in case of 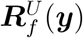 and ***J***^*U*^(***y***) being given.

However, the properties and the physical meanings of these quantifies 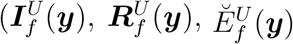, and 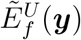 remain to be explored and clarified when a metric other than the potential energy is used as weight.

### 2.2 Computational details

Illustrative calculations have been performed in a model system, the alanine dipeptide in vacuum. The software package GROMACS (version 5.1.1)^47^ was employed for all-atom MD simulations with an integration time step of 0.5 fs. The same simulation setup as the ones in previous publications^27,28^ was used. Three non-equilibrium ensembles of short trajectories were collected: (1) TPE: The TPE for the transition from *C*_7eq_ to *C*_7ax_ harvested with transition path sampling, in which the definitions of two stable states (*C*_7eq_ and *C*_7ax_) were given based on the backbone dihedrals *ϕ* and *ψ*. Specifically, *C*_7eq_ corresponds to the phase space with −200° *< ϕ <* −55° (that is −180° *< ϕ <* −55° or 160° *< ϕ <* 180°) and −90° *< ψ <* 190° (that is −90° *< ψ <* 180° or −180° *< ψ <* −170°); *C*_7ax_ corresponds to the phase space with 50° *< ϕ <* 100° and −80° *< ψ <* 0°. The time length of each transition path is fixed to be 2 ps and the configurations were saved at every integration step. The obtained TPE consists of 5786 transition paths. (2) NPE_TS_: Ensemble of short trajectories shot from configurations near the transition state region, which was obtained with restrained MD simulations. The NPE_TS_ contains 5460 short trajectories with the time length of each trajectory being 2 ps. (3) NPE_TP_: Ensemble of short trajectories initiated from configurations taken from a small set of transition paths, which was taken from the TPE. It consists of 21260 short trajectories and the time length of each trajectory is 0.4 ps. NPE_TS_ and NPE_TP_ were considered as an approximation of the TPE and their trajectories were properly weighted according to the committor of configurations, from which these short trajectories were generated. These non-equilibrium ensembles have been previously used to test the viability of the EDARC approach and we refer the details of them to Ref. [^40^].

## 3 Results and Discussions

### 3.1 Free energy analogues are close approximations of free energy

We first calculated from the TPE the free energy analogue *Ă*^*U*^(***y***) defined in eq. 8 on the plane of *ϕ* and *θ*, which is shown in Fig. 1. One can identify two energy minimums, one around *ϕ* = −75 and *θ* = 0, and the other around *ϕ* = 60 and *θ* = 0. The energy barrier region or the transition state region is located around *ϕ* = 0 and *θ* = 0. The energy difference between the transition state region and the lowest energy minimum is about 22 kJ/mol.

**Figure 1:**
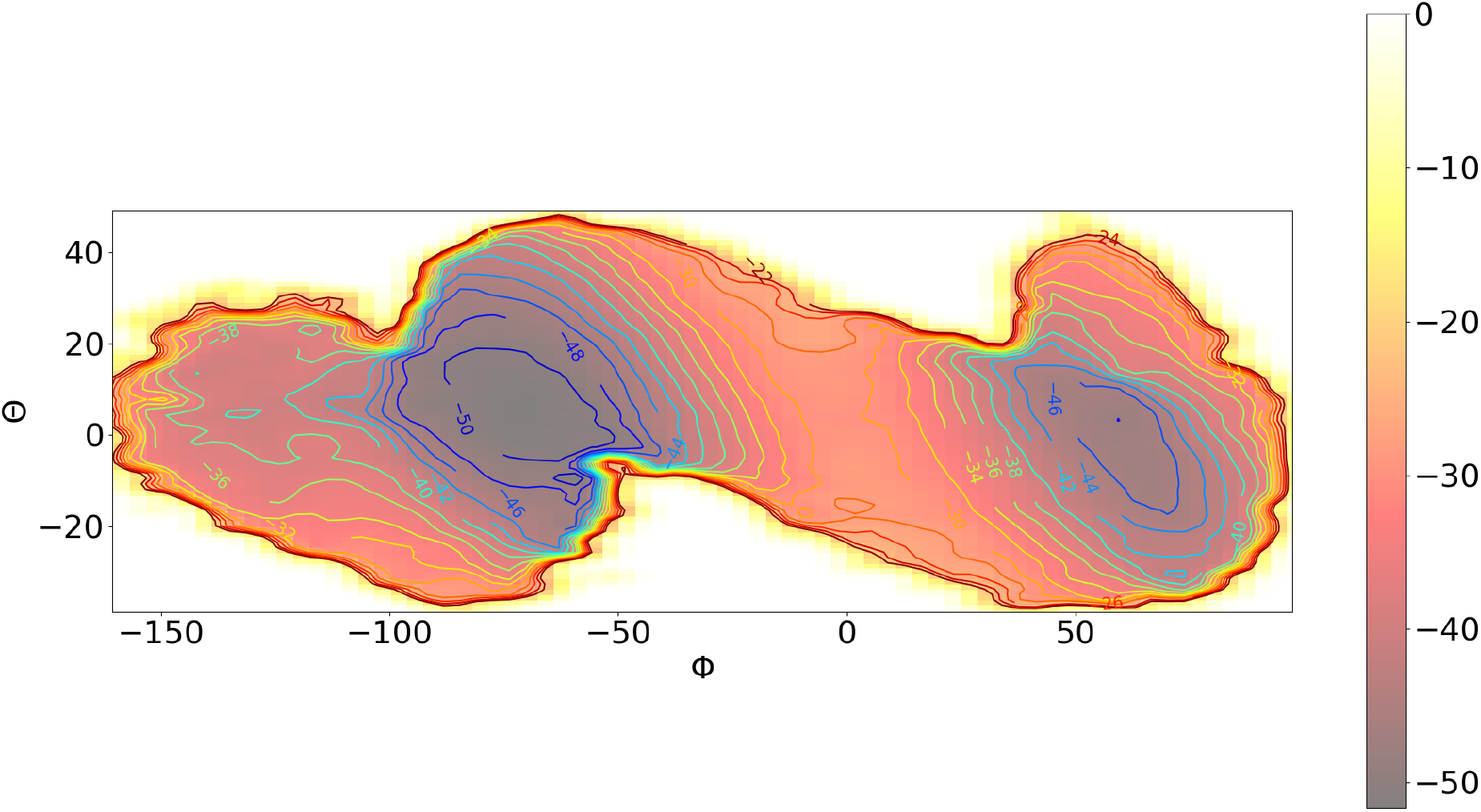
The free energy analogue *Ă*^*U*^(***y***) on the plane of *ϕ* and *θ*.

Then, we estimated the free energy analogue *Ã*^*U*^(***y***) from its gradients 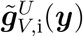 defined in eq. 18 (dashed contours in Fig. 2A) and compared it with *Ă*^*U*^(***y***) (solid contours in Fig. 2A). As indicated in eq. 18,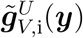 can be estimated only in the region where the reactive current is non-zero along both *ϕ* and *θ*. On the other hand, due to limited sampling, the uncertainty in the estimated 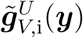 is very high in some regions, especially the periphery of the region that is sampled sufficiently (the well-sampled region), in which the reactive flux is not zero along any coordinate. Thus, the region where 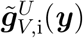 can be calculated in satisfying accuracy is largely not in a regular shape. As a result, the estimation of *Ã*^*U*^(***y***) from its gradients 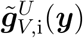 is numerically difficulty and the resulted *Ã*^*U*^(***y***) in the periphery of the well-sampled region looks strange. Nevertheless, the estimated *Ã*^*U*^(***y***) in the well-sampled region should be reliable. In the TPE, the reactive flux in the transition barrier region, the region between the reactant state and the product state, is expected to be divergence-free. We thus calculated the divergence of the reactive current in the plane of *ϕ* and *θ*, which is found to be vanishing small in the region *ϕ* ∈ [−30°, 30°] and *θ* ∈ [−25°, 25°] (Fig. S1). As shown in Fig. 2A, the free energy analogues *Ã*^*U*^(***y***) and *Ă*^*U*^(***y***) are largely consistent, especially in the transition barrier region where the reactive current is almost divergence-free. Thus, the result indicated in eq. 12 is confirmed in the plane of *ϕ* and *θ*.

**Figure 2:**
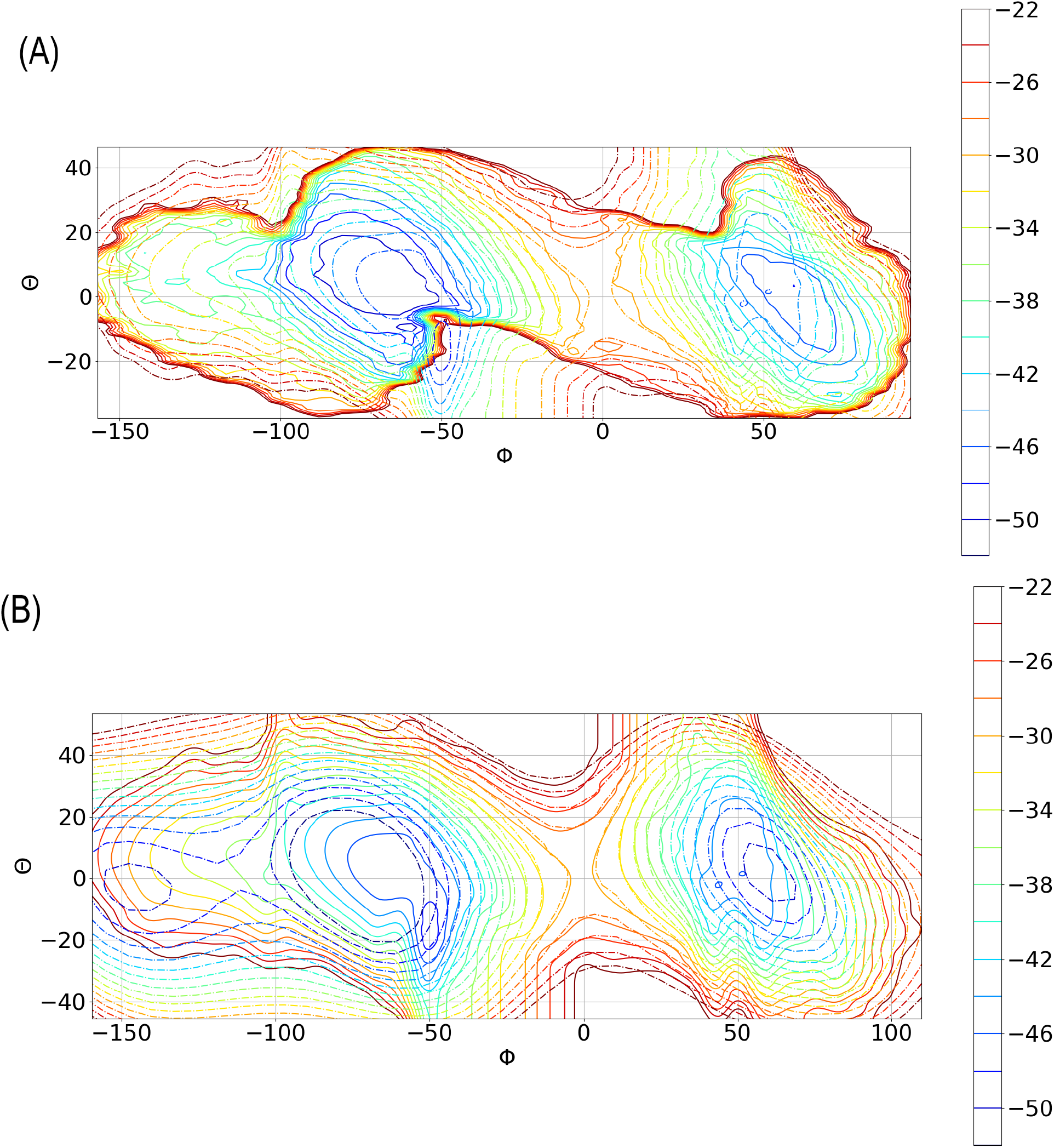
(A) Comparison between the free energy analogues *Ă*^*U*^(***y***) (solid contours) and *Ã*^*U*^(***y***) (dashed contours) on the plane of *ϕ* and *θ*. (B) Comparison between the free energy analogue *Ã*^*U*^(***y***) (solid contours) and the free energy (dashed contours) on the plane of *ϕ* and *θ*.

In the one-dimensional case, the free energy analogues *Ă*^*U*^(*ξ*) and *Ã*^*U*^(*ξ*) along the reaction coordinate are expected to be close approximations of the free energy. The free energy analogues *Ă*^*U*^(*ξ*) and *Ã*^*U*^(*ξ*) along an optimized reaction coordinate, a linear combination of *ϕ* and *θ*, were shown to be largely overlapped with the corresponding free energy in the transition barrier region.^40^ One would expect the free energy analogues *Ă*^*U*^(***y***) and *Ã*^*U*^(***y***) to be good approximations of the multidimensional free energy in the subspace of CVs as well. We thus calculated the free energy in the plane of *ϕ* and *θ* with umbrella sampling and compared it with *Ã*^*U*^(***y***) in Fig. 2B. Indeed, the contour map of *Ã*^*U*^(***y***) is overlapped to a great extent with the one of the free energy and they are almost identical in the region where the reactive current is shown to be divergence-free. Thus, the multidimensional free energy analogues proposed here are almost equivalent to the free energy. In case of that the TPE is more accessible than a long equilibrium trajectory, these free energy analogues can be evaluated as close approximations of the free energy to unveil important information on the reaction mechanism, which could be for instance the height of the free energy barrier and the locations of transition states, reactant states, and product states. Although multidimensional free energy can be evaluated by enhanced sampling methods such as umbrella sampling, these approaches usually require an informed guess of reaction coordinates. However, the TPE can be obtained with such path-sampling approaches as transition path sampling, which just requires order parameters to define metastable states. Thus, we believe that it is an advantage over traditional approaches to approximate the free energy with free energy analogues obtained from the TPE.

### 3.2 Reaction coordinate identification with 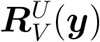

To explore the usefulness of 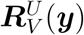 in identifying the reaction coordinate, we first calculated 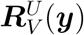 directly with eq. 10 (Fig. 3A). As highlighted in red boxes with dashed lines, there are three curves of 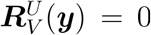, the left one is around *ϕ* = −75°, the one in the middle is around *ϕ* = 0°, and the right one is close to *ϕ* = −60°. These curves from left to right should correspond to the reactant state, the transition state, and the product state, respectively, as suggested in eq. 30. In the transition from the reactant state to the product state along the principal curve identified in Ref. [^30^] (blue dotted line in Fig. 3A), 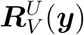 first increases from zero at the reactant state (around *ϕ* = −75° and *θ* = 7°) and reaches its maximum around *ϕ* = −42° and *θ* = 18°. It then decreases to its minimum near *ϕ* = 40° and *θ* = −10° and increases to zero at the product state (around *ϕ* = 60° and *θ* = 0°). Interestingly, the principal curve largely follows the ‘ridge’ of the contour map of 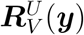 and passes characteristic points, i.e., the maximum, saddle points, and minimum, of 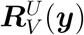. Thus, one should be able to identify from 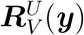 the principal curve and other important information of the reaction mechanism. As indicated in eq. 12, 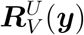 can also be calculated indirectly from the divergence of 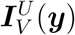, which can be obtained with eq. 7. As shown in Fig. 3B, the 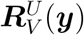 obtained with these two different ways are consistent in the region where the current is divergence-free, a result suggested by eq. 12.

**Figure 3:**
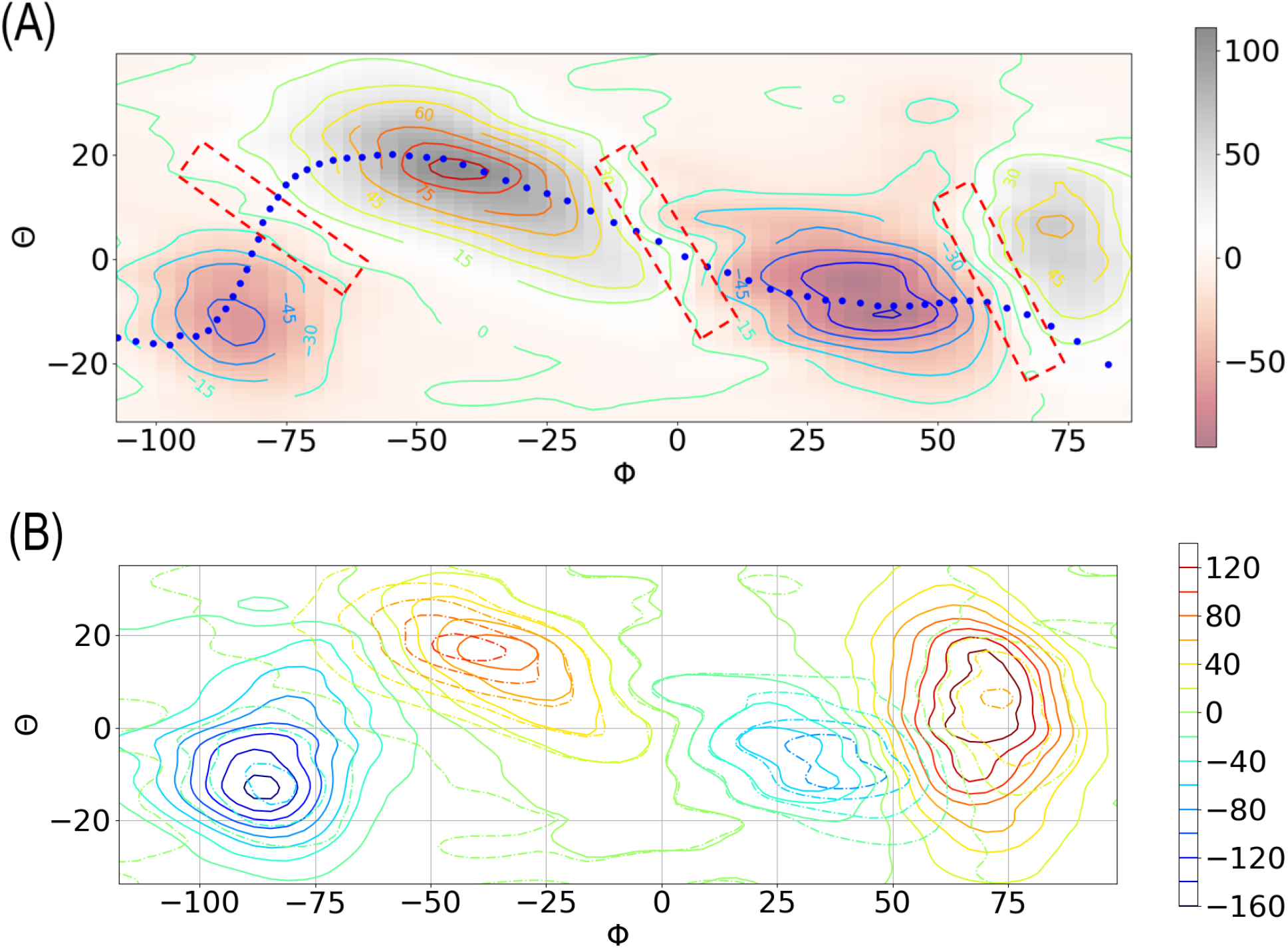
(A) The 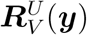 obtained directly by eq. 10 on the plane of *ϕ* and *θ*. The dotted blue line is the principal curve obtained in Ref. [^30^]. Three curves of 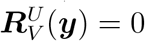 are highlighted in red boxes with dashed lines. (B) Comparison between the 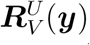 obtained directly by eq. 10 (dashed contours) and the one indirectly from the divergence of 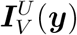 (solid contours) on the plane of *ϕ* and *θ*.

Importantly, we can estimate 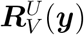 with eq. 16c, in which the potential energy components on the non-relevant coordinates are assumed to be negligible and thus are ignored. We thus calculated 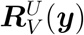 by taking only the potential energy components on *ψ* and *θ* into account, the resulted 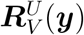 is shown in Fig. 4 and it is numerically more stable than the one obtained with eq. 10, in which the total potential energy is used instead (Fig. S2A). For example, the curves of 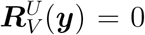 obtained with eq. 16c are almost linear, while the ones obtained with eq. 10 are fluctuating around the former ones (Fig. S2A). From the 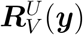 in Fig. 4, one can easily identify the stochastic separatrix, which corresponds to the curve of 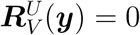 in the region around *ϕ* = 0°. Note that, the angle of the separatrix to the vector 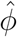 is about 120°. Thus the reaction coordinate, which is perpendicular to the separatrix, is about 30° to the vector 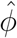, which is largely consistent with the ones identified in Ref. [^39^], where the committor information of all the configurations in the TPE were used. It is worth noting that the reaction coordinate is different to the eigenvector of the Hessian matrix of the free energy on this two-dimensional plane (Fig. 2), which is about 8° to the vector 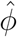.

**Figure 4:**
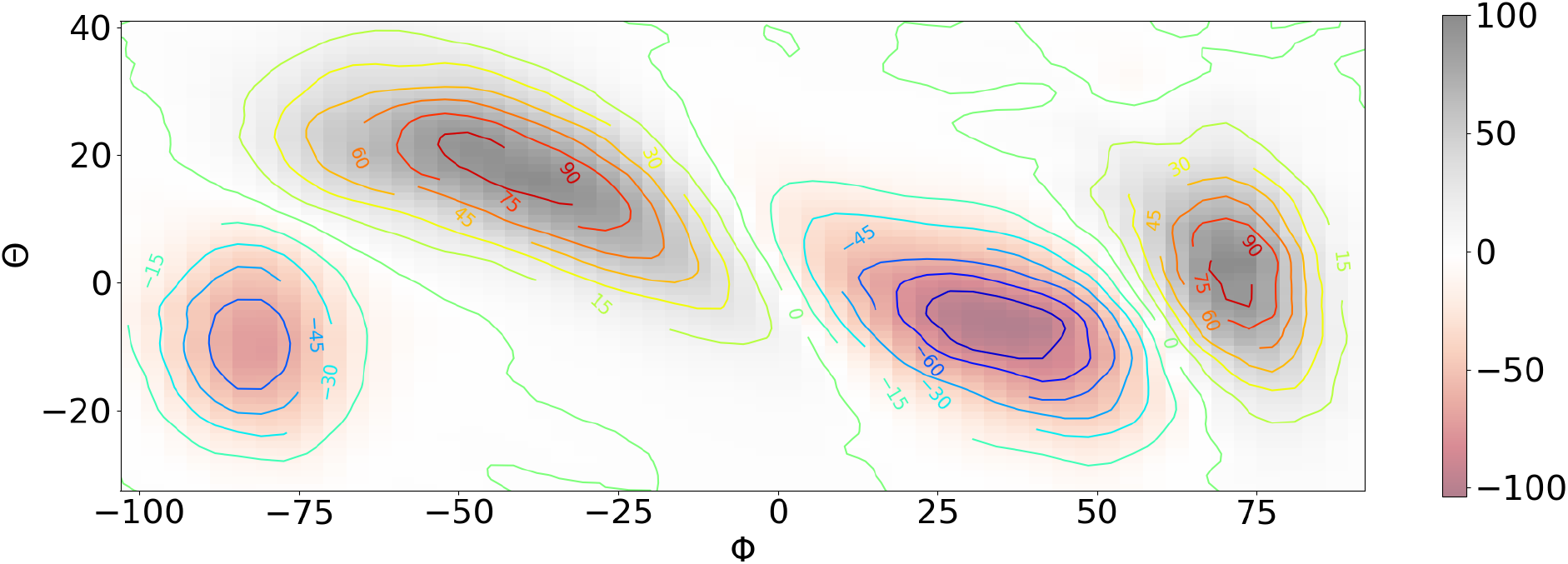
The 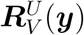 obtained directly by eq. 16c on the plane of *ϕ* and *θ*.

### 3.3 Evaluation the importance of a coordinate with 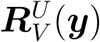

The components of 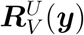 on each coordinate in the plane of *ϕ* and *θ* can also be used to quantify the energetic contribution of the coordinate to the transition from the reactant state to the product state. The 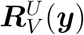 was thus further decomposed with eq. 13c into the components on BAT coordinates. We found that the components of 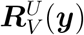 on *ϕ* and *θ* are the only two non-vanishing components (Fig. 5A and 5B), while the components on other coordinates are vanishingly small (Fig. S3). As shown in Fig. 5, the magnitude of the component on *ϕ* is about 4 times as much as the one on *θ*. Thus, the contribution from *ϕ* is thought to be about 4 times as much as the one from *θ*, which is consistent with the estimation obtained in the one-dimensional case with EDARC. ^40^ In the transition from the reactant state to the product state, the component of 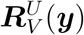 on *ϕ* is positive before crossing the separatrix and it is negative after crossing the separatrix, while the behaviour of the component on *θ* is the opposite. Such a result suggested that the contribution from *ϕ* promotes the transition, while the one from *θ* obstructs it, which is also consistent with the prediction from EDARC.^40^ In the component of 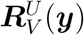 on *ϕ* in Fig. 5A, the curve of 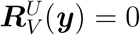 around the transition state is found to be perpendicular to the eigenvector of the Hessian matrix of the free energy and it suggests that the contribution from *ϕ* promotes the transition along the eigenvector of the Hessian matrix. In the component of 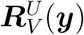 on *θ* in Fig. 5B, the curve of 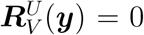 is nearly perpendicular to the separatrix and it indicates that the contribution from *θ* obstructs the transition along the reaction coordinate.

**Figure 5:**
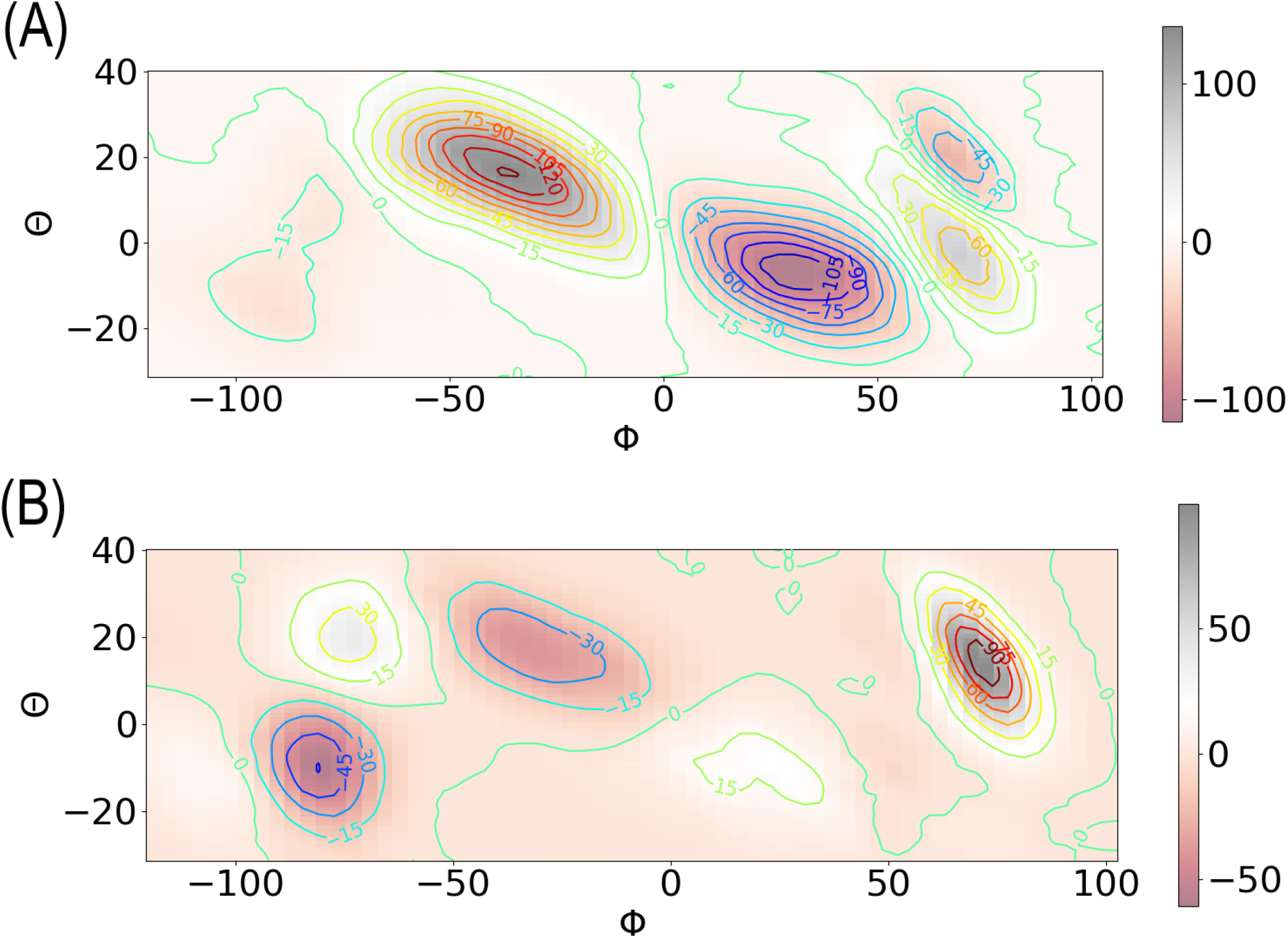
The components of 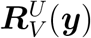 on *ϕ* (A) and *θ* (B) on the plane of *ϕ* and *θ*.

Usually, it is challenge to identify the best set of CVs to describe the dynamics of a transition process in complex systems. We thus ask whether 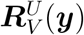 and its components can be used to quantify the importance of a coordinate to the transition on the the plane of *ϕ* and *ψ*, which is commonly considered as a set of CVs that is adequate but not as good as *ϕ* and *ψ* to describe the dynamics of alanine dipeptide in vacuum. Similar to the case in the plane of *ϕ* and *θ*, the 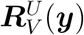 obtained both directly with eq. 10 and indirectly from the divergence of 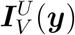 are consistent as well (Fig. S2B) in the transition barrier region (e.g., *ϕ* ∈ [−50°, 25°] and *ψ* ∈ [−80°, 40°]). A vector perpendicular to the line of 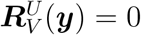 near *ϕ* = 0 is essentially in the *ϕ* direction, indicating that the role of *ψ* around the transition state region is negligible. In addition, as shown in Fig. S4, the components of 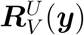 on *ϕ* and *θ* in this plane are largely non-zero while the ones on other coordinates are very small. Similar to the case on the plane of *ϕ* and *θ*, the magnitude of the component of 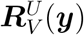 on *ϕ* is approximately 4 times as much as the one on *θ*. In addition, during the transition from the reactant state to the transition state, the contribution from *ϕ* is shown to be positive and the one from *θ* is negative. Thus, *ϕ* and *θ* can also be identified to be the most important coordinates for the *C*_7eq_ → *C*_7ax_ transition when a set of CVs with moderate quality is provided.

### 3.4 Results from non-equilibrium ensembles NPE_TS_ and NPE_TP_

Often-times, it is difficult to harvest the TPE for complex biomolecular systems, in which the time length of a transition trajectory can be in the scale of ns or even longer. We thus seek other non-equilibrium ensembles of short trajectories to best approximate the TPE. That is to say, the results of 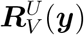 from other non-equilibrium ensembles should be consistent with the one from the TPE. We thus calculated the free energy analogue *Ã*^*U*^(***y***) from both NPE_TS_ and NPE_TP_ (Fig. S5), which were also shown to be consistent with the free energy, especially in the transition barrier region. With eq. 16c, we then calculated 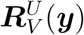 from NPE_TS_ and NPE_TP_, which are shown in Fig. 6A and 6B, respectively. Although the magnitude of 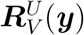 from these nonequilibrium ensembles (the TPE, NPE_TS_, and NPE_TP_) are different as the magnitude of the reactive current depends on the number of trajectories in the nonequilibrium ensemble, the contour maps of 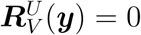 among these non-equilibrium ensembles are similar. Importantly, the curve of 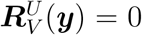 in the middle of the transition barrier region gives the separatrix, which is almost linear and is about 120° to the direction of 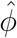. Thus, the separatrix in the plane of *ϕ* and *θ* identified from 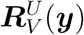 of both NPE_TS_ and NPE_TP_ is in well agreement with the one from the TPE. In addition, the magnitude of components of 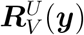 obtained from both NPE_TS_ (Fig. S6) and NPE_TP_ (Fig. S7) are mostly proportional to the ones from the TPE on the plane of *ϕ* and *θ* and they can also quantify the relative importance of a coordinate to the transition from *C*_7eq_ to *C*_7ax_. Thus, our approach can be applied to non-equilibrium ensembles other than the TPE and provides a good approximation of the free energy, the hypersurface of the separatrix, the relative importance of a coordinate to the transition process at any location of the subspace of CVs.

**Figure 6:**
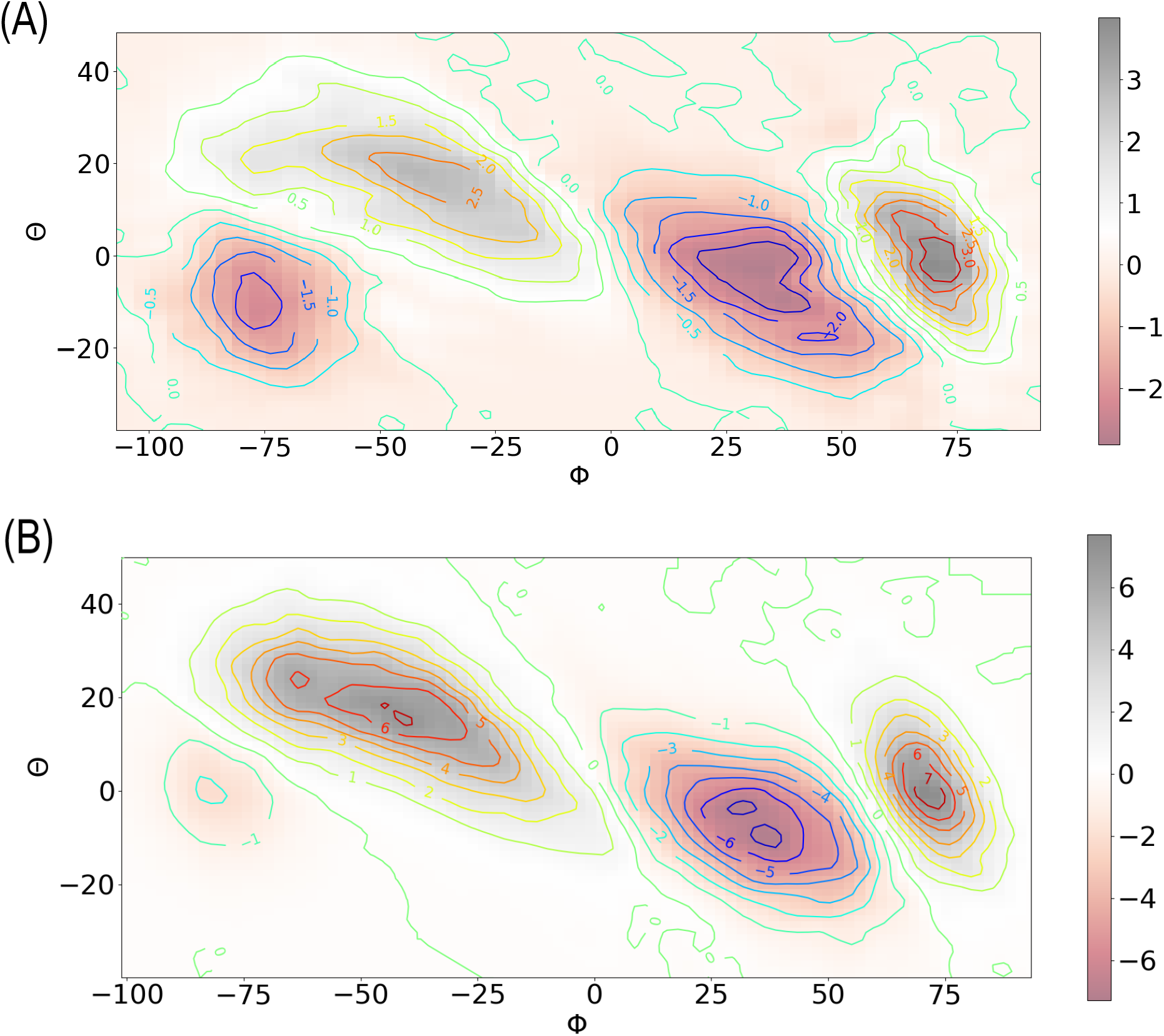
The 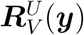 obtained directly by eq. 16c on the plane of *ϕ* and *θ* from non-equilibrium ensembles NPE_TS_ (A) and NPE_TP_ (B).

## 4 CONCLUDING REMARKS

We here proposed a theoretical framework for the potential energy weighted current and the total rate of change of potential energy. Upon projection of these two quantities to the subspace of CVs, which are assumed to capture the effective dynamics of a system, they can be used to obtain important information of the reaction mechanism, such as the reaction coordinate and the relative importance of a coordinate at any stage of a transition process. In the illustrative application to alanine dipeptide in vacuum, such a theoretical approach is demonstrated to be capable of reproducing the right one-dimensional representation of reaction coordinates and the contributions to the *C*_7eq_ → *C*_7ax_ transition at the per-coordinate level. Importantly, consistent results can be obtained from various non-equilibrium ensemble, some of which are easily available for complex biomolecular systems. We thus anticipate wide applications of the approach to biological and biochemical studies with MD simulations.

From the potential energy weighted current and TRCPE, free energy analogues *Ã*^*U*^(***y***) and *Ă*^*U*^(***y***) are then defined, respectively and they can be considered as the generalization of the one-dimensional free energy analogue proposed in Ref. [^40^] to multi-dimensional cases. Although *Ã*^*U*^(***y***) and *Ă*^*U*^(***y***) are equivalent at regions where the reactive current is divergencefree, *Ã*^*U*^(***y***) should be in a closer relation than *Ă*^*U*^(***y***) with the free energy at regions where the divergence of the reactive current does not vanish. From a practical aspect, we would like to emphasize that eq. 8 provides a convenient way to obtain a reasonable estimation of the multidimensional free energy in the region where the reactive current is divergence-free, while eq. 18 can be generally applied to calculate the free energy surface from a non-equilibrium ensemble in which the reactive current is non-vanishing. This is the case at least in the illustrative applications to alanine dipeptide in vacuum, as indicated in Fig. 2 and Fig. S5.

Interestingly, 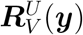 given in eq. 10 can be viewed as the directional derivative of the free energy analogue *Ã*^*U*^(***y***) and could be more useful than it appears. The curves of 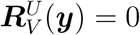 can provide the location of the separatrix, the reactant state and the product state. Along the principal curve, an approximation of the minimum free energy path, the direction of the reaction coordinate can be easily identified from the gradient of 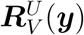 in the subspace of CVs and they are not necessarily the tangent of the principal curve.

We suggest that a few coordinates in close relevance to reaction coordinates can be first identified from flux maximization^28^ or one-dimensional free energy analogues with the EDARC approach,^40^ which do not require the calculation of partial derivative of potential energy with respect to coordinates (an independent and complete set), and then the relative importance of these relevant coordinates and a one-dimensional reaction coordinate as a (possibly curvilinear) function of them can be constructed by visual inspection of 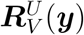 in subspaces of various combinations of CVs.

## 5 Appendix

### (A) The potential energy weighted current in the subspace of collective variables

The *i*-th element of 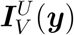 is the projection of 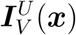 to *y*_i_ in the subspace of CVs, that is

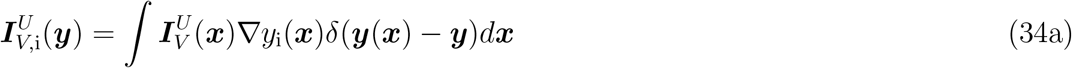

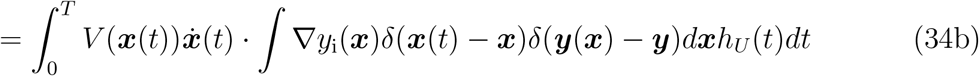

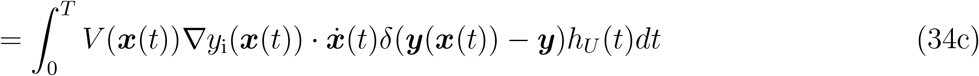

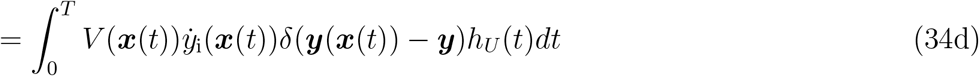

### (B) Proof of eq. 12

In the region of the subspace of the CVs where the divergence of the reactive current vanishes, i.e., ∇ · ***J***^*U*^(***y***) = 0, which indicates that there is no source or sink in this region for the ensemble *U*, we find that

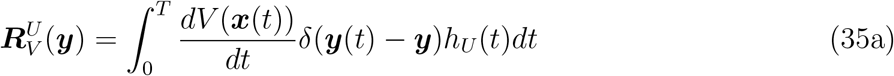

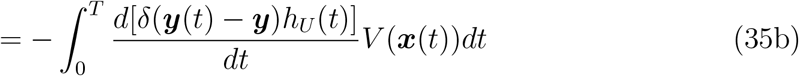

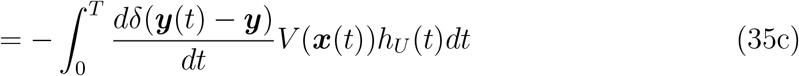

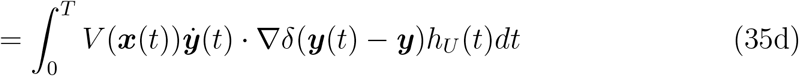

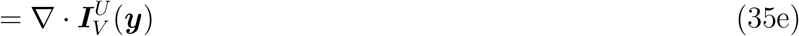

From eq. 35a to eq. 35b, integration by parts is applied. From eq. 35b to eq. 35c, we note that *dh*_*U*_ (*t*)*/dt* = 0 for any time when the trajectory hits the surface ***y***(***x***) = ***y***.

## Acknowledgement

This work was supported by Natural Science Foundation of Guangdong Province, China (Grant No. 2020A1515010984) and the Start-up Grant for Young Scientists (860-000002110384), Shenzhen University.

## Supporting Information Available

Divergence of the reactive current on the *ϕ* and *θ* plane; Comparison of 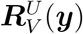 obtained with different equations; Components of 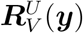 on the plane of *ϕ* and *θ* obtained from the TPE, NPE_TS_, and NPE_TP_; Components of 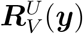 on the plane of *ϕ* and *ψ* obtained from the TPE; Comparison between *Ã*^*U*^(***y***) from NPE_TS_ or NPE_TP_ with the free energy.

## Notes

### Competing Interest Statement

The authors have declared no competing interest.

