## Supplemental Figures S1-S7 for "Potential Energy Weighted Reactive Flux and Total Rate of Change of Potential Energy: Theory and Illustrative Applications"

Wenjin Li
Institute for Advanced Study, Shenzhen University, Shenzhen, 518060, China

### Supplementary Material

#### Contents:

- Supplemental Figures S1-S7.

### Supplemental Figures

S1 Divergence of the reactive current on the $\phi$ and $\theta$ plane . . . . . . . . . . . . . . . . . . S2
S2 Comparison of $\boldsymbol{R}_V^U(\boldsymbol{y})$ obtained with different equations . . . . . . . . . . . . . . . . . . S3
S3 Components of $\boldsymbol{R}_V^U(\boldsymbol{y})$ on the plane of $\phi$ and $\theta$ (TPE) . . . . . . . . . . . . . . . . . . S4
S4 Components of $\boldsymbol{R}_V^U(\boldsymbol{y})$ on the plane of $\phi$ and $\psi$ (TPE) . . . . . . . . . . . . . . . . . . S5
S5 Comparison between $\tilde{A}^U(\boldsymbol{y})$ from $\mathrm{NPE_{TS}}$ or $\mathrm{NPE_{TP}}$ with the free energy . . . . . . . S6
S6 Components of $\boldsymbol{R}_V^U(\boldsymbol{y})$ on the plane of $\phi$ and $\theta$ ($\mathrm{NPE_{TS}}$) . . . . . . . . . . . . . . . . S7
S7 Components of $\boldsymbol{R}_V^U(\boldsymbol{y})$ on the plane of $\phi$ and $\theta$ ($\mathrm{NPE_{TP}}$) . . . . . . . . . . . . . . . . S8

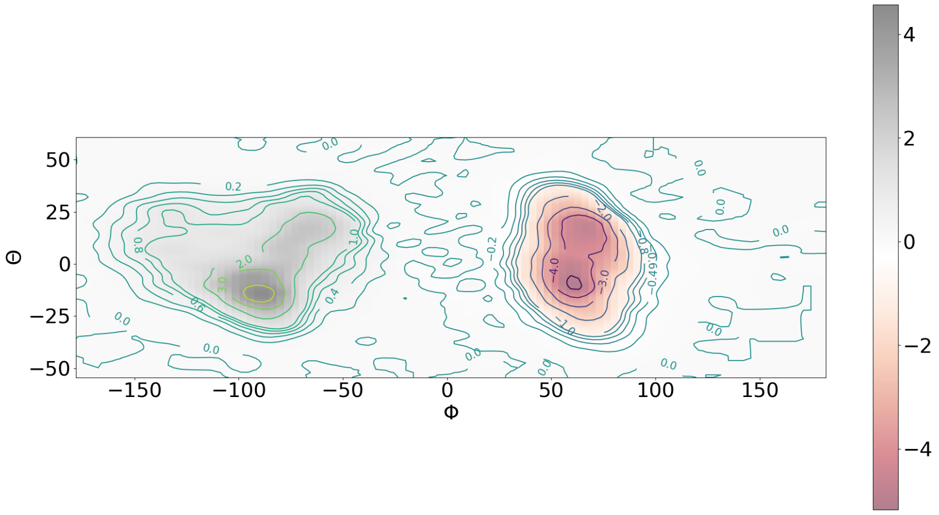


Figure S1: The divergence of the reactive current projected onto the $\phi$ and $\theta$ plane. The divergence of the reactive current at the region $\phi \in [-30°, 30°]$ and $\theta \in [-25°, 25°]$ is vanishingly small.

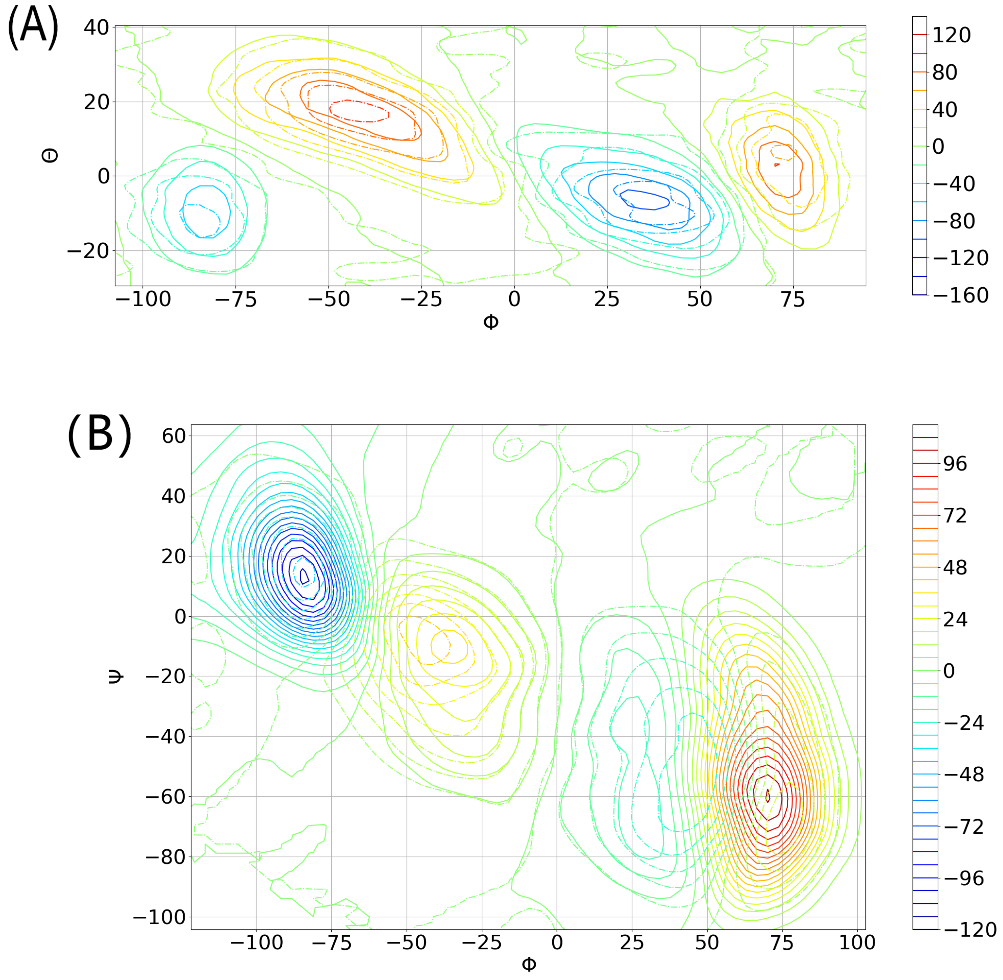


Figure S2: (A) Comparison between the $\boldsymbol{R}_V^U(\boldsymbol{y})$ obtained by eq. 16c (solid contours) and the one with eq. 10 (dashed contours) on the plane of $\phi$ and $\theta$. (B) Comparison between the $\boldsymbol{R}_V^U(\boldsymbol{y})$ obtained directly by eq. 10 (dashed contours) and the one indirectly from the divergence of $\boldsymbol{I}_V^U(\boldsymbol{y})$ (solid contours) on the plane of $\phi$ and $\psi$.

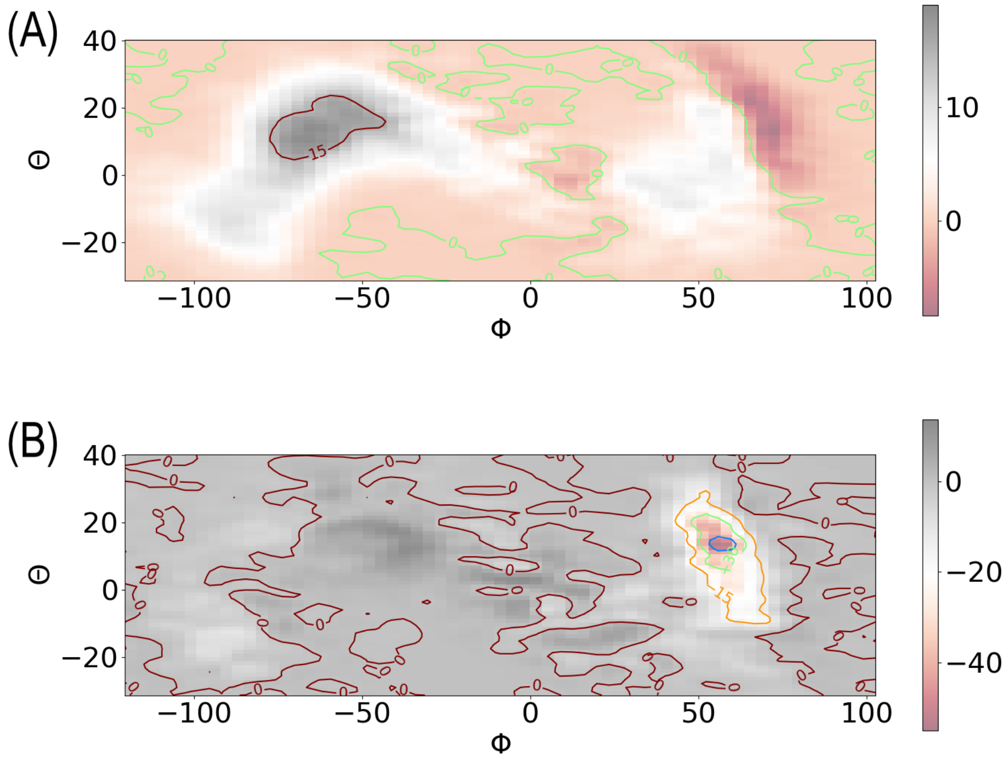


Figure S3: The components of $\boldsymbol{R}_V^U(\boldsymbol{y})$ on $\psi$ (A) and ang_N6_CA8_C14 (the 8th coordinate in the list of table S1 of Ref. [1]) (B) on the plane of $\phi$ and $\theta$.

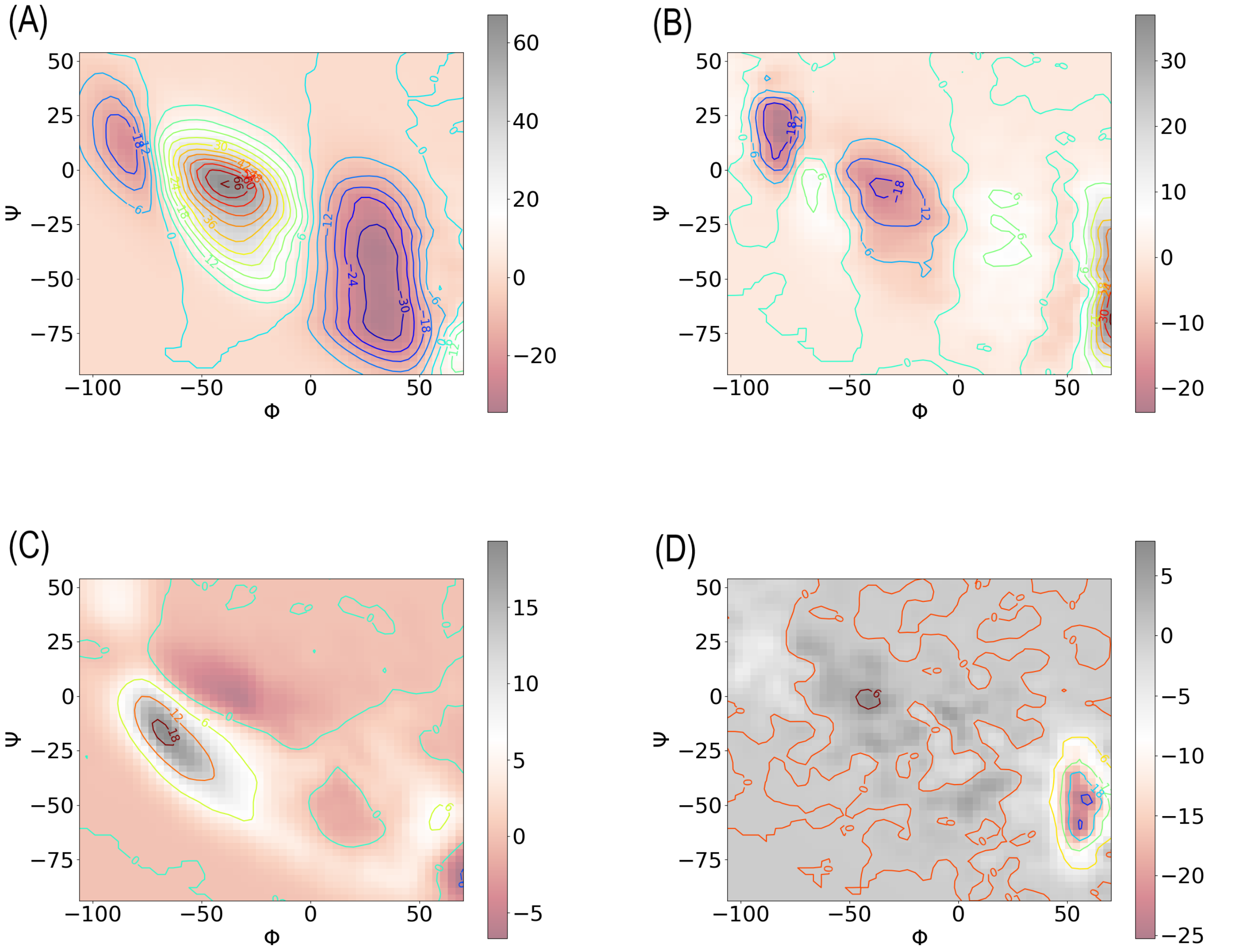


Figure S4: The components of $\boldsymbol{R}_V^U(\boldsymbol{y})$ on various coordinates on the plane of $\phi$ and $\psi$: (A) $\phi$, (B) $\theta$, (C) $\psi$, (D) ang_N6_CA8_C14.

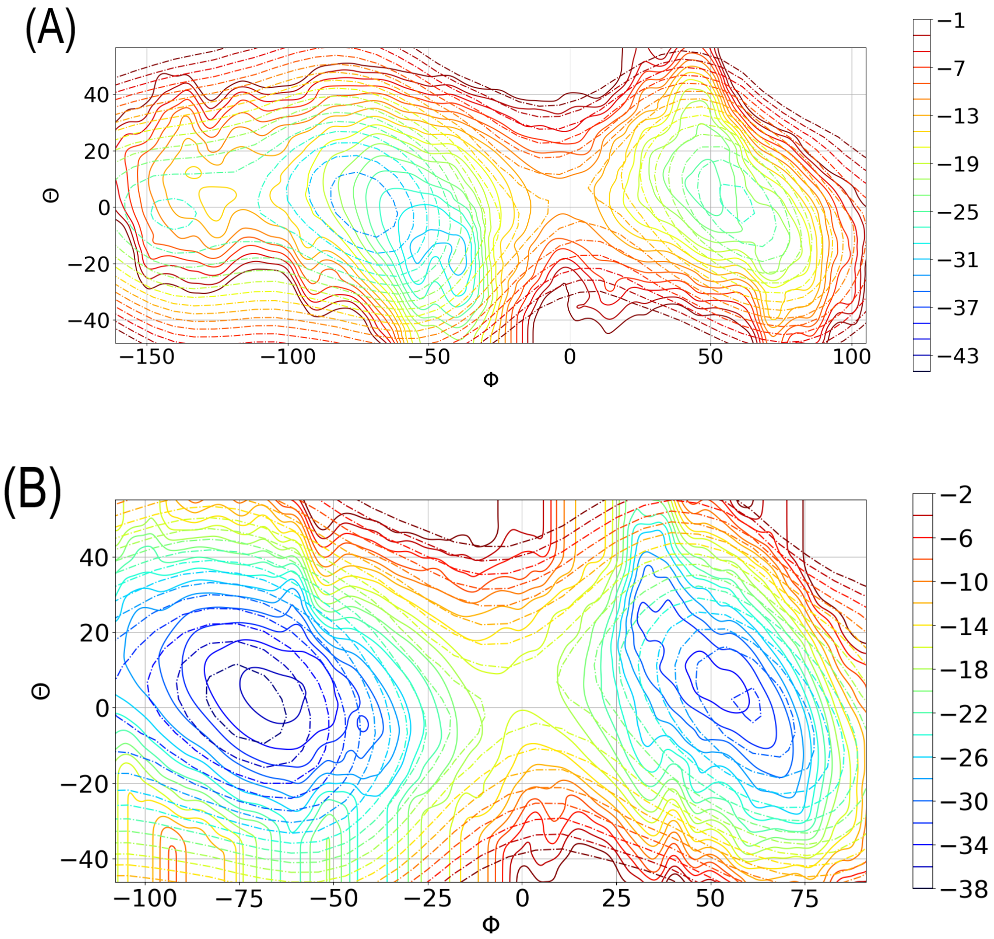


Figure S5: (A) Comparison between the free energy analogue $\tilde{A}^{U}(\boldsymbol{y})$ (solid contours) obtained from $\mathrm{NPE_{TS}}$ and the free energy (dashed contours) on the plane of $\phi$ and $\theta$. (B) Comparison between the free energy analogue $\tilde{A}^{U}(\boldsymbol{y})$ (solid contours) obtained from $\mathrm{NPE_{TP}}$ and the free energy (dashed contours) on the plane of $\phi$ and $\theta$.

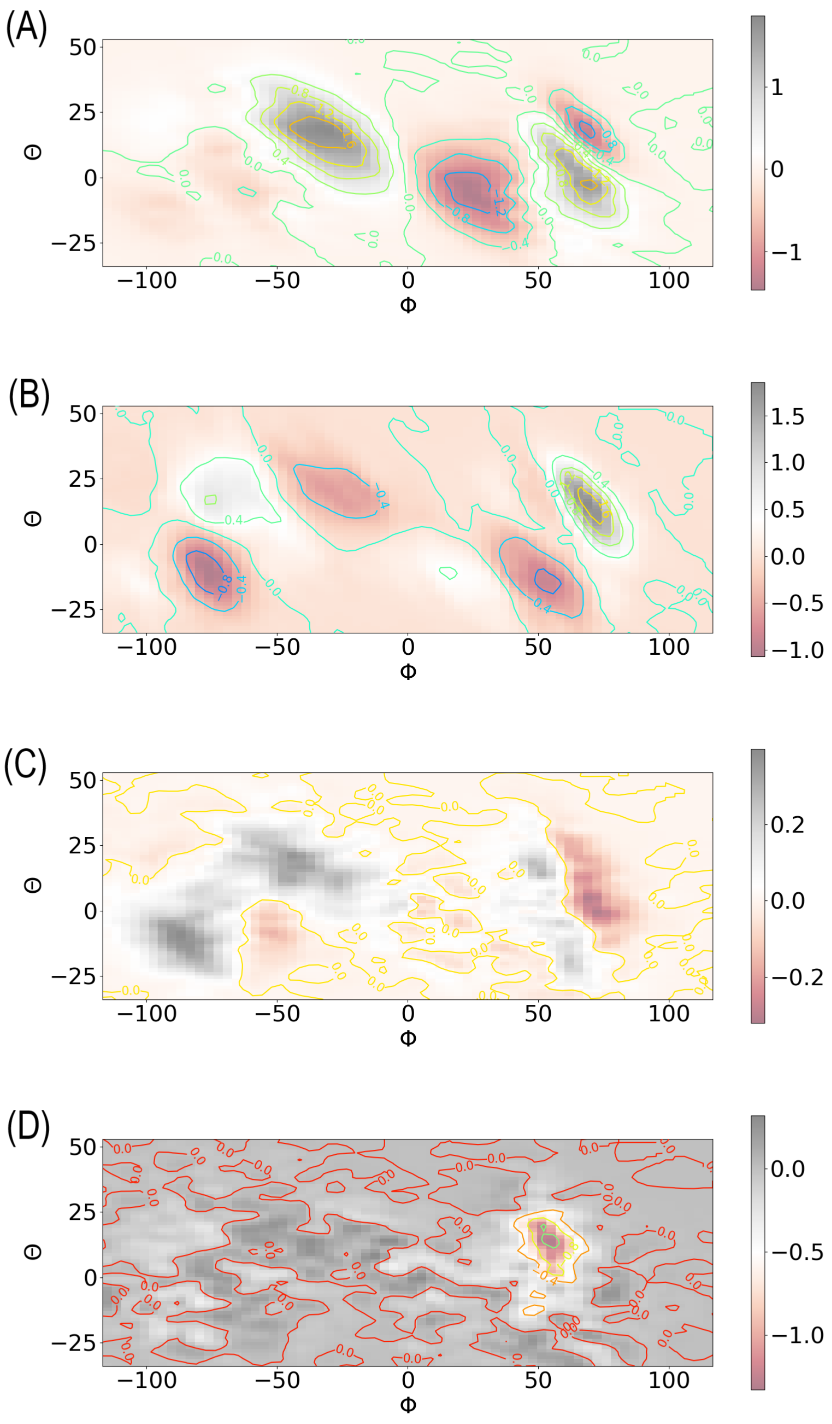


Figure S6: The components of $\boldsymbol{R}_V^U(\boldsymbol{y})$ on various coordinates on the plane of $\phi$ and $\theta$ from nonequilibrium ensemble $\mathrm{NPE_{TS}}$: (A) $\phi$, (B) $\theta$, (C) $\psi$, (D) ang_N6_CA8_C14.

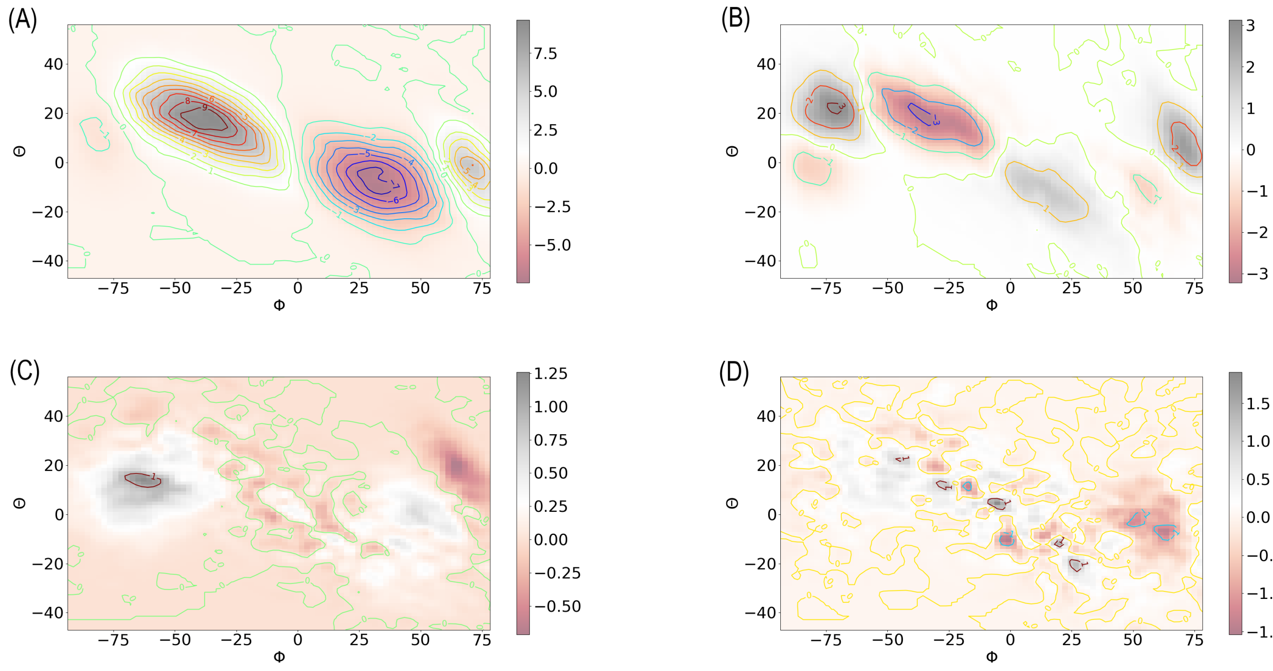


Figure S7: The components of $\boldsymbol{R}_V^U(\boldsymbol{y})$ on various coordinates on the plane of $\phi$ and $\theta$ from nonequilibrium ensemble $\mathrm{NPE_{TP}}$: (A) $\phi$, (B) $\theta$, (C) $\psi$, (D) ang_N6_CA8_C14.
